## Supplementary material for "BioCRNpyler: Compiling Chemical Reaction Networks from Biomolecular Parts in Diverse Contexts": S1 Text

### 1 Supplemental: Code for Examples

This section provides code from the synthetic biology examples figure (reproduced as Figure 1 in this document). The first three models are idealized in the sense that they are represented by Hill functions and include no cellular machinery such as ribosomes or polymerases. Producing these models in BioCRNpyler is easy and just requires the reuse of a few parts: `DNAassembly` represents a simple transcriptional unit with a promoter, transcript and optionally an ribosome binding site (RBS) and protein product. `RepressiblePromoter` creates a promoter modeled by a hill function. `Species` creates CRN species used in the models. Notice that only species which are shared between different `Components` need created by hand — BioCRNpyler takes care of the rest. For example, in inducible repression example the repressor $R$ is created by hand because it is placed inside the `RepressiblePrmoter`. However the DNA, transcript, and protein are automatically generated from the name of the `DNAassmebly`. Finally, everything is added together into a subclass of `Mixture` and compiled into a `ChemicalReactionNetwork`. The second three models build off the general architectures of the first three but add in more complicated context and implementation details. Instead of using `ExpressionDilutionMixture` and `SimpleTxTlDilutionMixture`, these models use the considerably more complex `TxTlDilutionMixture` which includes molecular machinery such as RNAP, ribosomes, RNAases, and background cellular processes. Additional implementation details in the form of `Components` and `Mechanisms` are also added to these models.

#### 1.1 Inducible Repression

Here a repressor is constituitively produces from a `DNAassembly`. This repressor is then linked to a `RepressiblePromoter` which models repression using a hill function.

```
from biocrnpyler import *
# Models a piece of DNA that constitutively produces the species R
repressor = Species("R")
const_rep = DNAassembly(name="const_rep", promoter="medium", rbs="medium",
     protein=repressor)
# R represses RepressiblePromoter which is placed into another DNAassembly reporter
prom = RepressiblePromoter(name="pR", repressor=repressor)
reporter = DNAassembly(name="Reporter", promoter=prom, rbs="strong",
     initial_concentration=1)
```

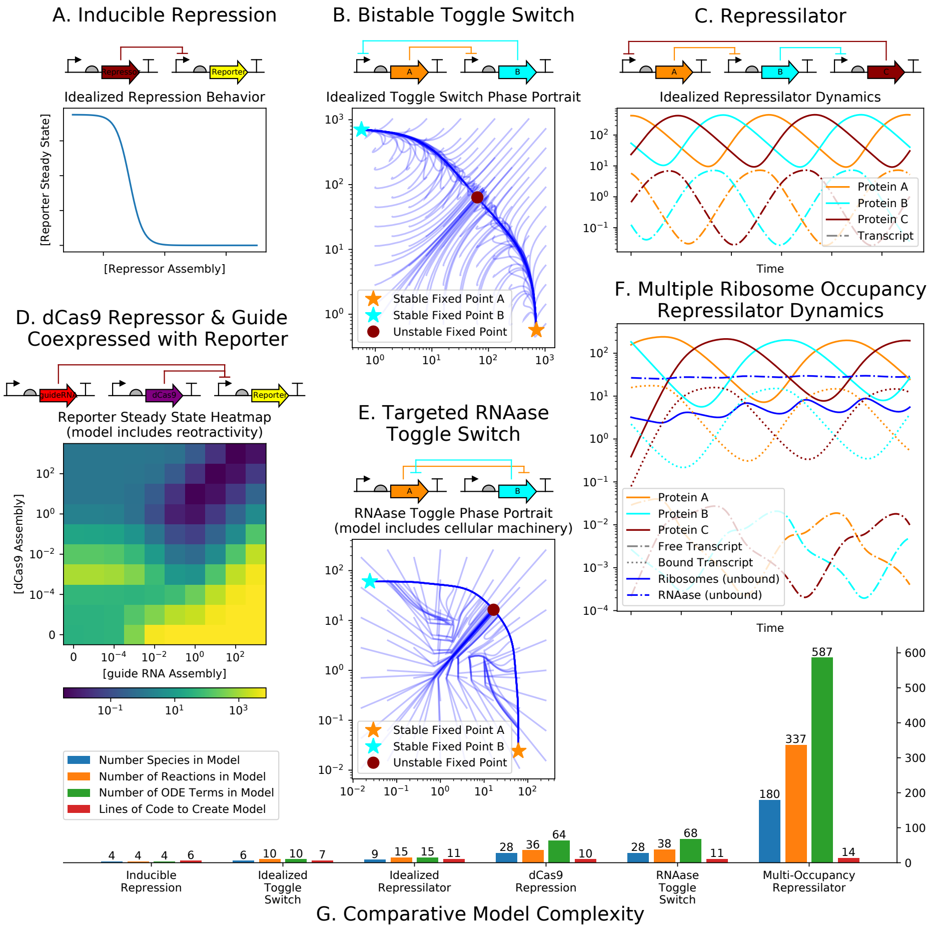


**Figure 1.** Motivating Examples. The idealized models (A, B, and C) do not model the cellular environment; genes and transcripts transcribe and translate catalytically. **A.** Schematic and simulation of a constituitively active repressor gene repressing a reporter. **B.** Schematic and simulations of of a toggle switch created by having two genes, $A$ and $B$, mutually repress each other. **C.** Schematic and dynamics of a 3-repressor oscillator. The detailed models (D, E, & F) model the cellular environment by including ribosomes, RNAases and background resource competition for cellular resources. **D.** A dCas9-guideRNA complex binds to the promoter of a reporter and inhibiting transcription. Heatmap shows retroactivity caused by varying the amount of dCas9 and guide-RNA expressed. The sharing of transcription and translational resources gives rise to increases and decreases of reporter even when there is very little repressor. **E.** A proposed model for a non-transcriptional toggle switch formed by homodimer-RNAase; the homodimer-RNAase made from subunit $A$ selectively degrades the mRNA producing subunit $B$ and visa-versa. **F.** A model of the Repressillator exploring the effects of multiple ribosomes binding to the same mRNA. **G.** Histogram comparing the sizes of models A-F and the amount of BioCRNpyler code needed to generate them.

```
# ExpressionDilutionMixture models gene expression without transcription/translation
mixture = ExpressionDilutionMixture(components=[reporter, const_rep],
     parameter_file="params.txt")
CRN = mixture.compile_crn()
```

#### 1.2 Toggle Switch

In the following example, a toggle switch is created by connecting two instances of `RepressiblePromoter` together. Notice that string names passed to promoter and rbs are used to help find parameters. BioCRNpyler comes with many default parameters to enable rapid model prototyping.

```
from biocrnpyler import *
# Creates A and is repressed by B
repA = Species("A")
promA = RepressiblePromoter(name="pA", repressor=repB)
assemblyA = DNAassembly(name="A", promoter=promA, rbs="medium", protein=repA,
     initial_concentration=1)
# Creates B and is repressed by A
repB = Species("B")
promB = RepressiblePromoter(name="pB", repressor=repA)
assemblyB = DNAassembly(name="B", promoter=promB, rbs="medium", protein=repB,
     initial_concentration=1)
# SimpleTxTlDilutionMixture includes transcription and translation but no machinery
mixture = SimpleTxTlDilutionMixture(components=[assemblyA, assemblyB], parameter_file=
     "params.txt")
CRN = mixture.compile_crn()
```

#### 1.3 Repressilator

The code to create a 3-node repession oscillator is really just adding one more unit and rewiring the toggle switch example.

```
from biocrnpyler import *
# Create Repressors
repA = Species("A")
repB = Species("B")
repC = Species("C")
# Create Promoters
promA = RepressiblePromoter(name="pA", repressor=repC)
promB = RepressiblePromoter(name="pB", repressor=repA)
promC = RepressiblePromoter(name="pC", repressor=repB)
#Create DNAassemblies
assemblyA = DNAassembly(name="A", promoter=promA, rbs="medium", protein=repA,
     initial_concentration=1)
assemblyB = DNAassembly(name="B", promoter=promB, rbs="medium", protein=repB,
     initial_concentration=1)
assemblyC = DNAassembly(name="C", promoter=promC, rbs="medium", protein=repC,
     initial_concentration=1)
# Place it all in a Mixture & Compile
mixture = SimpleTxTlDilutionMixture(components=[assemblyA, assemblyB, assemblyC],
     parameter_file="params.txt")
crn = mixture.compile_crn()
```

### 1.4 Cas9 Repressor and Guide RNA Coexpressed with Reporter

Modeling a dCas9-guideRNA repressor in bioCRNpyler requires that the dCAs9 and guide RNA know to bind together. This is accomplished via the `Component` subclass `ChemicalComplex` which models binding between multiple species. The resulting dCas9-guideRNA `ComplexSpecies` is used as a repressor.

```
from biocrnpyler import *
# parameter syntax: (mechanism_name, part_id, parameter_name) : value
# Only one dCas9-guideRNA complex binds to the promoter at once
params = {
    ("negativehill_transcription", None, "n"):1
}
# Create guide RNA and dCas9 Species
guide = Species("guide", material_type="rna")
dcas = Species("dCas9")
# These species will bind together by placing them in the ChemicalComplex Component
# "notdegradable" ensures that RNAases do not degrade gRNA-dCas9 complexes.
repressor = ChemicalComplex([dcas, guide], attributes=["notdegradable"])
reporter = Species("reporter")
# Constuitive Assemblies to produce dCas9 and the guideNRA
assembly_dcas = DNAassembly(name="dcas", promoter="medium", rbs="medium", protein=dcas)
assembly_guide = DNAassembly(name="guide", promoter="strong", rbs=None, transcript=guide)
# Create a repessible promoter
pReg = RepressiblePromoter(name="pA", repressor=repressor, parameters=params)
assembly_rep = DNAassembly(name="reporter", promoter=pReg, rbs="strong",
     protein=reporter, initial_concentration=1)
# Place the Components in a Mixture
extract = TxTlDilutionMixture("e coli", components=[assembly_rep, assembly_dcas,
     assembly_guide, repressor], parameter_file="params.txt")
#Compile the CRN
crn = extract.compile_crn()
```

### 1.5 Targeted RNAase Toggle Switch

The targeted RNAase toggle switch model is a hypothetical model similar to a normal toggle switch but with regulation at the RNA level instead of the transcriptional level. This is accomplished by creating two constitutively expressed RNAases (which are `ChemicalComplexes` made up of two subunits) and adding custom `Mechanisms` to the `Mixture` modeling the degradation of any species with the attribute “tagA” and “tagB” by RNAase A and RNAase B, respectively.

```
#Create an RNA species with degradation tag sequence tagB
TA = Species("A", attributes=["tagB"], material_type="rna")
#Create homodimer subunit A
A = Species("A", material_type="protein")
#RNAase A is a homodimer made up of two identical subunits
rnaaseA = ChemicalComplex([A]*2)
#create a DNAassembly that produces A
assemblyA = DNAassembly(name="A", promoter="strong", transcript=TA, rbs="medium",
     protein=A, initial_concentration=1)

#Same as above but for Species B
TB = Species("B", attributes=["tagA"], material_type="rna")
B = Species("B", material_type="protein")
rnaaseB = ChemicalComplex([B]*2)
assemblyB = DNAassembly(name="B", promoter="strong", transcript=TB, rbs="medium",
     protein=B, initial_concentration=1)
#add all the Components to a Mixture
mixture = TxTlDilutionMixture("e coli", components=[assemblyA, assemblyB, rnaaseA,
```

```
    rnaaseB], parameter_file="default_parameters.txt")
# Deg_Tagged_Degredation Mechanism makes rnaaseA degrade anything with attribute "tagA"
mixture.add_mechanism(Deg_Tagged_Degredation(mechanism_type="tagA_degredation",
    deg_tag="tagA", protease=rnaaseA.get_species()))
# Deg_Tagged_Degredation Mechanism makes rnaaseB degrade anything with attribute "tagB"
mixture.add_mechanism(Deg_Tagged_Degredation(mechanism_type="tagB_degredation",
    deg_tag="tagB", protease=rnaaseB.get_species()))
#Compile the CRN
CRN = mixture.compile_crn()
```

### 1.6 Multiple Ribosome Occupancy Repressilator Dynamics

Simulating multiple-ribosome occupancy in the Repressilator mostly reuses the code from Section 1.3 with the main addition of a new `Mechanism` to model transcription being placed into the `Mixture`.

```
#Create Repressors
repA = Species("A")
repB = Species("B")
repC = Species("C")
#Create Promoters
promA = RepressiblePromoter(name="pA", repressor=repC)
promB = RepressiblePromoter(name="pB", repressor=repA)
promC = RepressiblePromoter(name="pC", repressor=repB)
#Create DNAassemblies
assemblyA = DNAassembly(name="A", promoter=promA, rbs="strong", protein=repA,
    initial_concentration=1)
assemblyB = DNAassembly(name="B", promoter=promB, rbs="strong", protein=repB,
    initial_concentration=1)
assemblyC = DNAassembly(name="C", promoter=promC, rbs="strong", protein=repC,
    initial_concentration=1)
#Extra parameters for the Multi_tx Mechanism
extra_params = {"max_occ":10, ("multi_tx", None, "k_iso"):50, ("multi_tl", None,
    "k_iso"):50, "cooperativity":2}
#Add Everything to a Mixture
mixture = TxTlDilutionMixture("e coli", components=[assemblyA, assemblyB, assemblyC],
                            parameter_file="default_parameters.txt",
                                parameters=extra_params, overwrite_parameters=True)
#Add the multi_tl translation mechanism to the mixture, overwriting the old one.
mixture.add_mechanism(multi_tl(name="multi_tl",
    ribosome=mixture.ribosome.get_species()), overwrite=True)
#Compile the CRN
crn = mixture.compile_crn()
```

### 1.7 Lac Operon Model

The Lac Operon model combines described in the main text (with the accompanying figure reproduced here as Figure 2) contains many of the elements of the previous examples together into a considerably more complex circuit. First, the lacR monomer can be inhibited by binding to allolactose, producing a `ChemicalComplex`. These monomers can bind to eachother (either with or without allolactose), resulting in 3 possible dimers, also modeled as `ChemicalComplexes`. Finally, pairs of these possible dimers can bind together combinatorially to produce 6 forms of the lacR tetramer, modeled as 6 `ChemicalComplexes`. The regulatory protein CRP binding to camp is also modeled as a `ChemicalComplex`, denoted c-CRP. Combinatoric binding reactions of all these different transcription factors is modeled using a powerful `Component` called a `CombinatorialConformationPromoter`. Briefly, this component makes use of a `PolymerConformation`; a special kind of species abstracted as a hypergraph. Nodes of

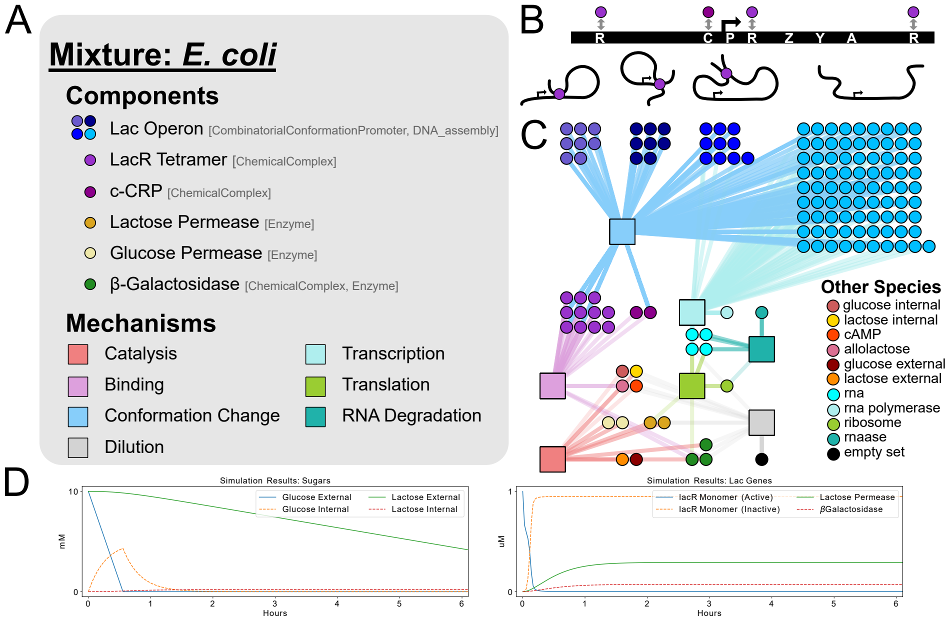


**Figure 2.** A model of the lac operon compiled using BioCRNpyler specifications with 141 species and 271 reactions using ∼50 lines of code. **A.** A `Mixture` contains a set of `Components` and `Mechanisms`. The `Component` classes used for each element of the model are shown in brackets. **B.** A schematic of the lac operon and the three looped and one open conformation it can take. Each conformation contains a combinatoric number of states based upon the accessible binding sites: R are lac repressor binding sites; C is the activator c-CRP binding site; P is the promoter; and Z, Y, A are the three lac genes. **C.** A graph representation of the compiled CRN. Each circle is a unique chemical species. Square boxes show how chemical species interact via reactions generated by specific `Mechanisms`. **D.** Simulated output of the model.

this graph are species, including monomers inside of one or more polymers. Hyper-edges of this graph are multisets of nodes bound together.

A `CombinatorialConformationPromoter Component` enumerates all possible binding reactions from a set of initial states through a set of intermediate states, to a set of final states with the constraint that species can only be added to the polymer between each set of states. Here, each state is a user defined `PolymerConformation`. This allows for all the possible binding pathways resulting in some fully "bound" or saturated polymer state (including loops) to be easily enumerated one binding event at a time. Moreover, specific hyper-edges of the `PolymerConformation` can be set to make the `CombinatorialConformationPromoter` active or inactive to model transcriptional regulation (or, alternatively, be modeled using custom rates for each conformation). Transport of glucose and lactose into the cell are modeled using `Enzymes` to represent glucose permease and lactose permease. Finally, metabolism of lactose by $\beta$-galactosidase is modeled using two `Components`: an `Enzyme` and `ChemicalComplex` (because $\beta$-galactosidase is a homeotetramer). In total, this results in 16 `Components` which are placed in a `Mixture`.

```
#LacR Tetramer can have multiple isoforms
lacR_m = Species("LRm", attributes = ["machinery"]) #lac R monomer
allolactose = Species("allolactose") #allolactose
lacR_m_i = ChemicalComplex([lacR_m, allolactose]) #Inhibited lacR monomer
lacR_d = ChemicalComplex(2*[lacR_m]) #LacR dimer (active)
lacR_d_i = ChemicalComplex([lacR_m, lacR_m_i]) #lacR dimer (inactve)
lacR_d_2i = ChemicalComplex([lacR_m_i, lacR_m_i]) #lacR dimer (inactve)
lacR_t = ChemicalComplex(2*[lacR_d.get_species()]) #LacR tetramer
lacR_t_i = ChemicalComplex([lacR_d.get_species(), lacR_d_i.get_species()]) #tetramer
    with 1 inhibited dimer
lacR_t_2i = ChemicalComplex([lacR_d.get_species(), lacR_d_2i.get_species()]) #tetramer
    with one double inhibited dimers
lacR_t_2ib = ChemicalComplex(2*[lacR_d_i.get_species()]) #tetramer with two inhibited
    dimers
lacR_t_3i = ChemicalComplex([lacR_d_i.get_species(), lacR_d_2i.get_species()]) #tetramer
    with two inhibited dimers
lacR_t_4i = ChemicalComplex(2*[lacR_d_2i.get_species()]) #tetramer with two double
    inhibited dimers



camp = Species("camp")
CRP = Species("CRP", attributes = ["machinery"])
cCRP = ChemicalComplex([camp, CRP], attributes = ["machinery"])

glu_ex, glu_in = Species("glucose", attributes = ["external"]), Species("glucose",
    attributes = ["internal"])
lac_ex, lac_in = Species("lactose", attributes = ["external"]), Species("lactose",
    attributes = ["internal"])
lacPermease = Enzyme("lacPermease", substrates = [lac_ex], products = [lac_in])

gluPermease = Enzyme("GluPermease", substrates = [glu_ex], products = [glu_in, camp],
    attributes = ["machinery"])

betaGalm = Species("betagal_monomer") #BetaGal Monomer
betaGal_c = ChemicalComplex(4*[betaGalm])
betaGal_e = Enzyme(enzyme = betaGal_c.get_species(), substrates = [lac_in], products =
    [allolactose])

#Binding sites of the lac operon
L = Species("L") #LacR binding site
C = Species("C") #camp-CRP binding site
P = Species("P") #Promoter

#Combine these binding sites into a PolymerConformation
```

```python
O = PolymerConformation(polymer = [L, C, P, L, L]) #Unbound conformation

#These lists will be used to enumerate the different functional states of the lac operon
inactivating_complexes = [] #List of complexes which stop transcription
single_bound_conformations = [] #list of conformations with a single site bound
loop_conformations = [] #A list of PolymerConformations with loops
loop_complexes = [] #A list of the ChemicalComplexes forming the loops

#Iterate through lacR binding sites
for i in [0, 3, 4]:
    single_bound_conformations.append(Complex([O.polymers[0][i],
         lacR_t.get_species()]).parent)
    single_bound_conformations.append(Complex([O.polymers[0][i],
         lacR_t_i.get_species()]).parent)
    single_bound_conformations.append(Complex([O.polymers[0][i],
         lacR_t_2i.get_species()]).parent)

    if i == 3:
        inactivating_complexes.append(Complex([O.polymers[0][i], lacR_t.get_species()]))
        inactivating_complexes.append(Complex([O.polymers[0][i], lacR_t_i.get_species()]))
        inactivating_complexes.append(Complex([O.polymers[0][i],
             lacR_t_2i.get_species()]))

single_bound_conformations.append(Complex([O.polymers[0][1], cCRP.get_species()]).parent)

#Create Loops from single bound complexes
for lind, (i, j) in enumerate([(3, 4), (0, 4), (0, 3)]):
    CO1 = Complex([single_bound_conformations[lind*3].polymers[0][i],
         single_bound_conformations[lind*3].polymers[0][j], lacR_t.get_species()])
    CO2 = Complex([single_bound_conformations[lind*3+1].polymers[0][i],
         single_bound_conformations[lind*3+1].polymers[0][j], lacR_t.get_species()])
    CO3 = Complex([single_bound_conformations[lind*3+2].polymers[0][i],
         single_bound_conformations[lind*3+2].polymers[0][j], lacR_t.get_species()])
    #loop_complexes.append(CO1)
    if i == 3 or j == 3:
        inactivating_complexes.append(CO1)
    loop_conformations += [CO1.parent, CO2.parent, CO3.parent]

#Bind cCRP to each loop the complex
saturated_loop_conformations = []
for LO in loop_conformations:
    saturated_loop_conformations.append(Complex([LO.polymers[0][1],
         cCRP.get_species()]).parent)

#the Lac operon is modeled as a Promoter and a DNAassembly
CCP = CombinatorialConformationPromoter(
    intermediate_states = single_bound_conformations,
    final_states = saturated_loop_conformations,
    promoter_location = 2,
    promoter_states = [],
    promoter_states_on = False, #This will make all states ON by default
    inactivating_complexes = inactivating_complexes, #[loop_complexes[0],
         loop_complexes[2]], #Any state containing either of these complexes will be OFF
    activating_complexes = []
)

A = DNAassembly(name = "lac", dna = O, promoter = CCP, rbs = "strong", protein =
     [betaGalm, lacPermease], initial_concentration = 5*10**-2)

#Machinery in E. Coli
rnap = Species("RNAP", attributes = ["machinery"], material_type = "protein")
ribosome = Species("Ribo", attributes = ["machinery"], material_type = "protein")
rnaase = Species("RNAase", attributes = ["machinery"], material_type = "protein")

#Create TxTl Mechansisms
mech_tx = Transcription_MM(rnap = rnap)
```

```
mech_tl = Translation_MM(ribosome = ribosome)
mech_cat = MichaelisMenten()
mech_bind = One_Step_Binding()

#Create Global Dilution Mechanisms
#all dna-species, machinery species, and external species are not diluted.
mech_dil = Dilution(
    filter_dict = {"dna":False, "machinery":False, "conformation":False,
         "external":False, "lacPermease":True},
    default_on = True
)
mech_rna_deg = Degredation_mRNA_MM(nuclease = rnaase)

#Create a Mixture
mechanisms = [mech_tx, mech_tl, mech_cat, mech_bind, mech_rna_deg, mech_dil]
components = [lacR_m_i, lacR_d, lacR_d_i, lacR_d_2i, lacR_t, lacR_t_i, lacR_t_2i,
     lacR_t_2ib, lacR_t_3i, lacR_t_4i, cCRP, A, lacPermease, gluPermease, betaGal_c,
     betaGal_e]

#set some parameters manually
params = {
    "kf":100.0, "kr":10.0, "kcat":1.0
}

M = Mixture(
    name = "e coli",
    parameters = params, parameter_file = "default_parameters.txt",
    components = components,
    mechanisms = mechanisms
)

#Choose initial concentrations
initial_conc = {
    camp:0,
    CRP:1.0,
    lacR_m:1.,
    glu_ex:10000.,
    lac_ex:10000.,
    gluPermease.get_species():5.
}
#Compile the CRN
CRN = M.compile_crn(initial_concentration_dict = initial_conc)
```

### 1.8 Component Enumeration Examples

Component enumeration (both local and global) allows for `Components` and `Mixtures` to generate additional `Components` using arbitrary Python code. Importantly, by generating additional `Components`, this process preserves the downstream functions of the BioCRNpyler compilation algorithm – in short allowing enumerated `Components` to generate species and reactions via `Mechanisms`. Custom `ComponentEnumerators` can be easily made by subclassing either the `LocalComponentEnumerator` or `GlobalComponentEnumerator` classes and defining the subclasses' `enumerate_components(component, keywords)` function which returns a list of `Components`. For a `LocalComponentEnumerator` the first argument "component" will be a single `Component`. For a `GlobalComponentEnumerator` the first argument "component" will be a list of `Components`. To highlight the power of component enumeration, BioCRNpyler currently has two `ComponentEnumerators` designed to compile sophisticated synthetic biological circuits. These are described and illustrated in main text with the component enumeration figure recreated here as Figure 3. In this section, we provide code and more elaborate explanations of the different models.

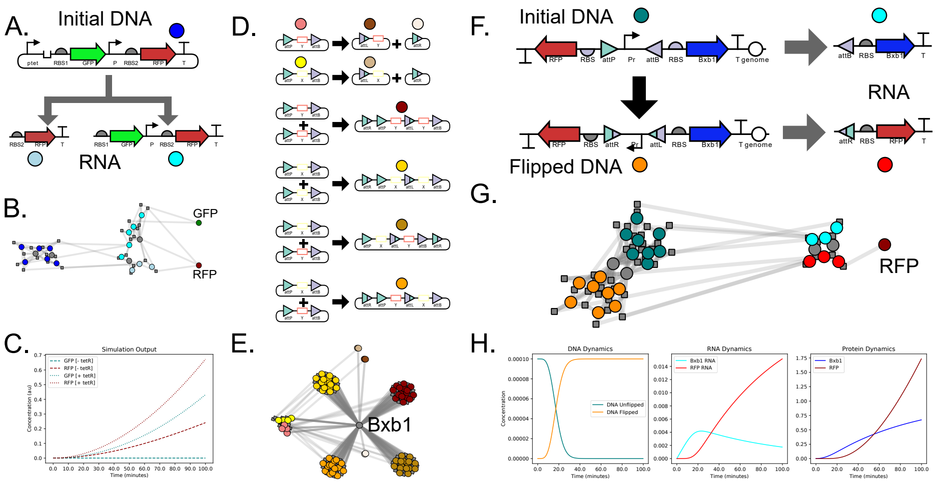


**Figure 3.** Examples involving component enumeration. A. Schematic of local component enumeration for a gene expression circuit where a single DNA `Component` generates multiple RNA `Components`. B. The CRN for (A) represented graphically. Colored dots are species corresponding to the components adjacent to the dots in (A). C. Simulated output from the CRN in (B). D. Schematic of global component enumeration in an integrase circuit where one or more DNA `Components` recombine to produce new DNA `Components`. Note that the larger DNA outputs could also recombined analogously but this is not shown. E. The CRN for (D) represented graphically. Colored dots are species which correspond to the components adjacent to the dots in (D). F. A genetic circuit which combines global and local component enumeration to flip a promoter which drives gene expression. G. The CRN for the circuit in (F). Colored dots are species representing the components adjacent to the dots in (F). H. Simulated output of the CRN.

#### 1.8.1 DNA Construct Local Component Enumeration Example

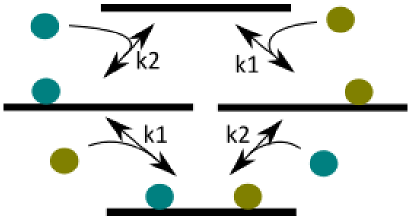


**Figure 4.** Two independent binding reactions. Two species (colored circles) can bind to a piece of DNA (black line) independently. Each species must be able to react with unbound DNA and also bound DNA. Because the binding is independent, the rate constant for binding is unchanged whether each species reacts with a bound or unbound DNA.

When dealing with simple DNA species that contain only a single promoter and produce only a single RNA, reactions like the binding or RNA polymerase (RNAP) to the promoter can be easily generated when a user creates an promoter `Component`. The goal of `DNA_construct`, however, is to allow an arbitrary ordering and assemblage of parts. Thus it should be possible for a single piece of DNA to contain many promoters acting independently. This presents a computational problem to BioCRNpyler in that a variety of bound species must be automatically created. If two separate DNA molecules existed with independent promoters, it would be sufficient to simply populate binding and unbinding reactions independently. However, if we imagine both promoters present on the same piece of DNA, it would be possible to have a states (represented as species) where with RNA polymerase bound in either, both, or neither promoter as illustrated in Figure 4. We will refer to this as *combinatorial binding*. BioCRNpyer solves the problem of combinatorial binding, as well as parsing the intricate behavior of arbitrary orders of `Promoters`, `RBSes` (ribosome binding sites), `Terminators`, etc. using a `LocalComponentEnumerator` inside the `DNA_construct Component`.

Combinatorial binding occurs when at least two sites on one species can be bound independently by different species. The identity of the species which performs the binding is not important; the only thing that matters is that it is possible for any of the independent binding sites to be occupied and none of the sites is affected by any of the other ones. Our goal in implementing combinatorial binding in BioCRNpyler was to re-use the existing interface for DNA parts like `Promoters`, but allow them to be used with complicated DNA species that can combinatorially multiple promoters. Another consequence of combinatorial binding is that it is essential to keep track of which binding site is occupied. For example, in the condition where two promoters exist on one DNA, it is important to know which of them is bound by RNA polymerase in the case that only one promoter is bound, because that determines which RNA will be produced.

For this reason we have created the `OrderedPolymer` class, which serves as a way to organize species with multiple binding sites. An `OrderedPolymer` is a list where each element has a reference to its location within the list and also a link back to the parent. This is important because of the interaction with `Components` such as `Promoters`. Since a `Promoter` must accept a DNA species as input, now we can use an element from the `OrderedPolymer` class (called an `OrderedMonomer`) as an input. Thus, a `Promoter` can use this `OrderedMonomer` in exactly the same way as a standard DNA species would have been used, but now any `ComplexSpecies` created as a result of `Mechanisms` contained within that `Promoter` will be created in the proper binding position of the parent `OrderedPolymer`. Likewise all the parts contained in a `DNA_construct` are also stored in an `OrderedPolymer`. Simply using an `OrderedPolymer` does not accomplish

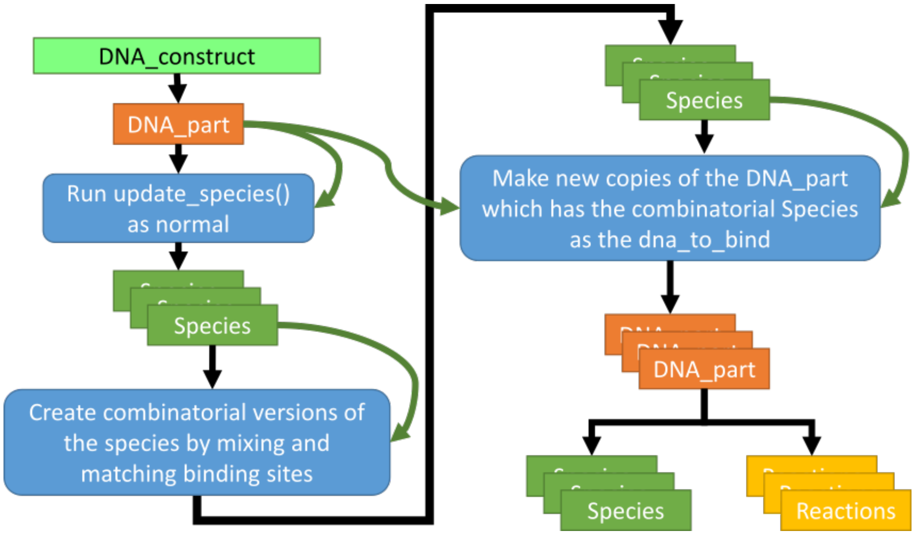


**Figure 5.** Combinatorial Enumeration. The function `update_species()` is called twice in combinatorial enumeration. First, each `DNA_part` is asked to `update_species` while binding to an "empty" version of the `DNA_construct` underlying species. This generates a set of possible bound states with each active `DNA_part` in isolation (first stack of green "Species" boxes). Then, these `ComplexSpecies` are recombined: for example, a complex at position 1 can be combined on the same molecule with a complex on position 2 (second stack of green "Species" boxes). Once these combinatorial complexes are made, a copy of each active `DNA_part` is made, each containing a different combinatorial complex. Finally, these copied `DNA_parts` are made to `update_species()` and `update_reactions()`, thus generating all the proper species and reactions in between the combinatorially recombined versions of the original `DNA_construct`. Black arrows represent logical flow, and green arrows represent information flow. For example, information from the `DNA_parts` present in the original `DNA_construct` is combined with the new species generated from the combinatorial enumeration to produce the copied `DNA_parts` which are essential for producing the proper species and `Reactions`.

combinatorial binding. Given a `DNA_construct` that contains multiple active `Components`, we allow each `Component` to generate the bound species relevant to its `Mechanisms`, then create all combinatorial combinations of these bound species. This means, for example, if Promoter 1 yields a `ComplexSpecies` where RNAP binds to position 1, and Promoter 2 yields a `ComplexSpecies` where RNAP binds to position 2, there is a possible combinatorial `ComplexSpecies` where both RNAPs are bound. Once this combinatorial species is generated, it can be fed back into the active `DNA_parts` (`Components` are also `DNA_parts`) in order to generate the proper reactions and species. The logical flow of this process is illustrated in Figure 5.

Given that all the `Components` contained within a `DNA_construct` have had all relevant variables properly populated, this process will always yield the correct species and reactions for a given `DNA_construct`. A different algorithm, which we call `TxTl_explorer`, is used to populate `Components` properly, according to the "central dogma". This method iterates sequentially along the linear `DNA_construct` in order, tabulating all `Components` that exist between `Promoters` and `Terminators` and thus end up contained in RNAs, as well as `CDS` components which exist following `RBS` components. `TxTl_explorer` operates in the forward and reverse directions, as well as being able to cross from one side of a `DNA_construct` back to the beginning in case we are trying to represent a circular plasmid.

Within each Mixture, multiple DNA_constructs may be present. Combinatorial enumeration is able to properly create the necessary species and reactions pertaining to a single DNA_construct, and the DNA_parts contained within. A necessary consequence of combinatorial enumeration, however, is the creation of RNA_constructs. These are similar linear arrangements of parts except now an RBS fills the role of a Promoter. Thus, during CRN compilation, DNA_constructs lead to the creation of RNA_constructs. Once species and reactions are created for all DNA_constructs in the mixture and all RNA_constructs that started life in the mixture as well as those that are generated, nothing else needs to be done. Thus we call this combinatorial enumeration ”local” because each DNA_construct only knows about itself, and there is no potential for infinite recursion. This is in contrast with integrases, as presented in the next section.

The following code shows the DNA_construct created in Figure 3A which uses local component enumeration to figure out all the possible bound conformations of the species and generate the correct RNA_constructs.

```
#TetR repressor species
tetR = Species("tetR")

#Create DNA parts

#Promoters
P1 = RegulatedPromoter("ptet",[tetR],leak=False) #repressed by tetR
P2 = Promoter("P") #constiuitive

#Ribosome binding sites
rbs1 = RBS("RBS1")
rbs2 = RBS("RBS2")

#Coding sequences
cGFP = CDS("GFP")
cRFP = CDS("RFP")

#Terminator
T = Terminator("T")

#Combine the parts into a DNA Construct (plasmid)
#the DNA_construct automatically creates its own Local Component Enumerator
plasmid = DNA_construct(
    [P1, rbs1, cGFP, P2, rbs2, cRFP, T],
    initial_conc = 0.0001, circular = True
)

#create an extra parameter to control integration rate
parameters = {"kint":0.05}

#Create a Mixture
M = TxTlExtract(
    name = "e coli extract", components = [plasmid],
    parameters = parameters, parameter_file = "default_parameters.txt"
)

#compile the CRN
CRN = M.compile_crn()
```

### 1.9 Integrase Global Component Enumeration Example

Integrases are capable of rearranging DNA based on the identity and layout of DNA sequences called attachment sites. The mechanism by which different biochemical integrases undertake their reactions can be different, but the outcome is usually similar: a recombination event occurs between two attachment sites. In the case of serine integrases such as Bxb1, two different sites known as attB and attP react to form attL and attR, recombining the DNA in the process. From the point of view of what the recombination actually does to the sequences being recombined, five basic types of integration are possible, as illustrated in Figure 6.

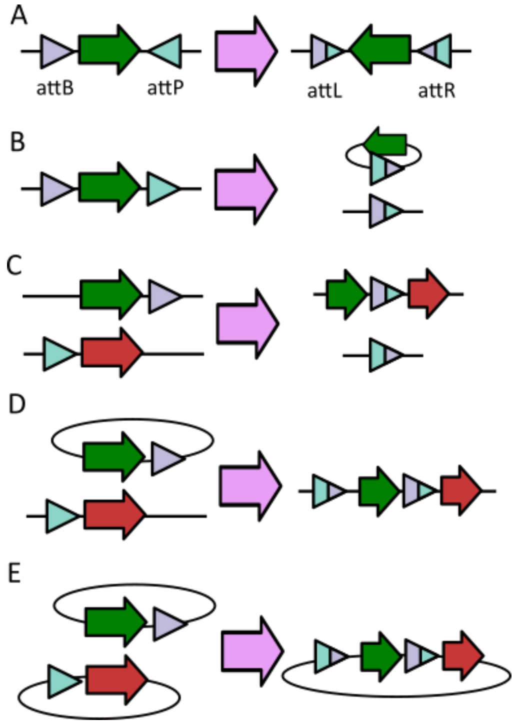


**Figure 6.** Topology changes induced by integrase. Triangles represent integrase sites, black line represents DNA, thick arrow illustrates the direction of the DNA. (A) Flipping: integrase sites on the same piece of DNA are pointed at each other and the DNA between the integrase sites is inverted in direction. (B) Deletion: integrase sites on the same piece of DNA are pointed in the same direction, and the DNA between the integrase sites is excised into a circular fragment. (C) Integration between linear DNA: integrase sites are present on two different pieces of DNA, which are both linear. This results in two pieces of DNA containing the recombined sites, that are also both linear. (D) Integration of a circular DNA: integrase sites are present on two different pieces of DNA, but one of them is circular. This results in one linear piece of DNA containing the recombined pieces. (E) Integration of two circular DNAs: Integrase sites are present on two different pieces of DNA, and both are circular. This results in one circular piece of DNA with essentially the same organization as D.

Since `DNA_constructs` represent linear sequences of parts, the transformations they undergo as a result of integrase activity are fairly straightforward, and lead to the creation of new `DNA_constructs`. These new `DNA_constructs` might have different RNA or protein products, owing to the fact that promoters might now be arranged differently with regard to terminators or protein coding genes might now be oriented in forward or reverse directions, and these eventualities are handled by local enumeration, as presented in the previous section. The challenge to BioCRNpyler, and ultimately to the user, is when to stop generating these new constructs? One might consider a simple case such as that presented in Figure 6A. Integrase sites facing each other lead to a portion of DNA being "flipped". Due to the nature of serine integrases, the reaction does not proceed in the reverse direction, so it seems like generating a single new `DNA_construct` with the proper sequences flipped should be the end.

However, in a real cell there could be multiple copies of the same DNA sequence, whether that DNA exists as a plasmid or a cell is replicating and thus has multiple genomes, or the cell was transformed with many linear copies of the same DNA. Thus integrase could perform a recombination reaction between attB and attP on two different pieces of DNA. We call this type of reaction "intermolecular" to distinguish it from the "intramolecular" flip reaction which would occur between two sites present on the same piece of DNA. Thus even a simple flip reaction can lead to infinite possible `DNA_constructs`, and for this reason we have included a variety of flags that allow users to decide whether intermolecular reactions are allowed, and sequentially how many times the pool of `DNA_constructs` in a mixture are interrogated for new `DNA_constructs`.

Once BioCRNpyler has enumerated all the possible integrase reactions that can occur, it is important to make sure that the resulting CRN includes the proper reactions to connect together the generation and destruction of these new `DNA_constructs`. This is more sophisticated than normal reaction and species generation for a couple of reasons. First, the species that are involved with integrase reactions are in general already generated. Reactant and product `DNA_constructs` have already been generated as part of the first step which is enumerating possible integrase reactions. When an integrase recombines a piece of DNA, we assume that the intervening DNA goes along for the ride, and any complexes and bound proteins present there should not be affected by the action of the integrase. This means, for example, that if a promoter is to be flipped, then any polymerase that happened to be bound to that promoter could get flipped as well. Thus, the correct CRN should contain a reaction going from unbound DNA to unbound, flipped DNA, but also from RNAP-bound DNA to RNAP-bound flipped DNA.

As we mentioned above, in local component enumeration, a copy is made of each `DNA_part` which contains a species representing a different bound state of the rest of the DNA molecule. Integrase sites are likewise copied, but in a way integrase sites are not independent as all other `DNA_parts` are assumed to be, since they care about the state of their partner site. In the case of an intermolecular reaction, an integrase site must care about all the different combinatorially bound states of the other `DNA_construct` that it will integrate with, and create the proper reactions. For this reason we call this step "global enumeration", and the process is a bit more complicated.

As indicated in Figure 7, a recursive step allows `DNA_constructs` to create new `DNA_constructs` using the logic embedded in an integrase-specific `GlobalComponentEnumerator`. As part of this process, the integrase reactions (flip, deletion, integration, etc) which will take place are recorded within `IntegraseSites` which are a type of `DNA_part`. Then, during local component enumeration, these saved integrase reactions are used to perform integrase reactions on species containing other bound species, which were generated as part of local component enumeration. These recombined species are used to create reactions that involve integrase. No new species are generated using the integrase `GlobalComponentEnumerator`, as the integrase binding to integrase sites has already been taken care of during local component enumeration. Real integrases have an intermediate step where a tetramer of integrase molecules creates a bridge between the two attachment sites, and this type of reaction would be a good improvement to add in future work.

The recursion inherent in global component enumeration reveals that even relatively simple integrase-containing constructs can lead to very complicated populations of different DNA sequences inside cells. It may be possible to ignore this complexity, as in the flipping example it is likely that intramolecular recombination is much more

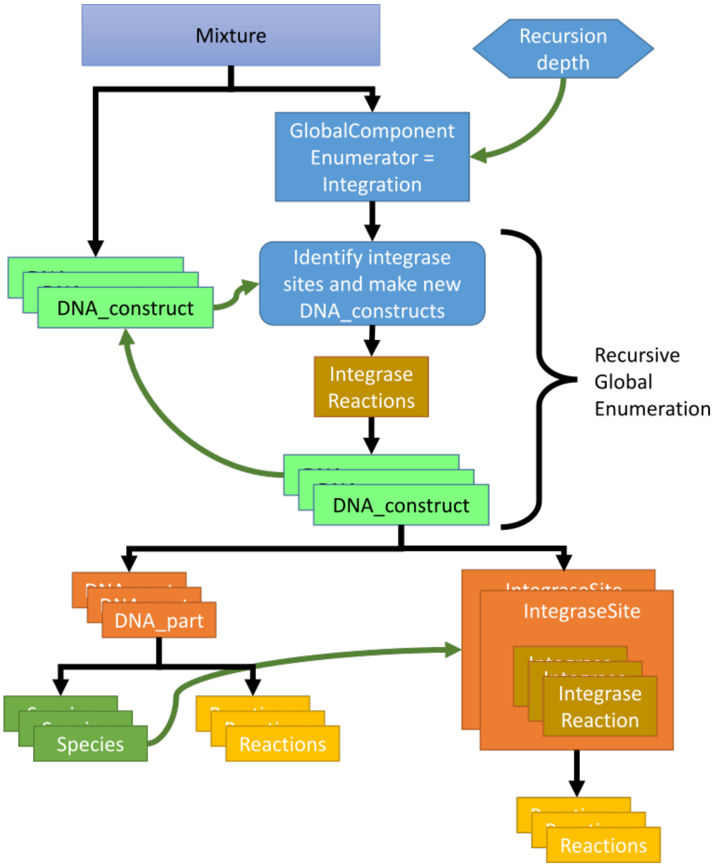


**Figure 7.** Global enumeration. A mixture (blue box at the top) starts by containing `DNA_constructs` and a `GlobalComponentEnumerator`. A user-entered parameter of recursion depth is fed into the `GlobalComponentEnumerator` to limit how many times new `DNA_constructs` will be generated. The global enumeration loop is indicated by the curly brackets. All the `DNA_constructs` in the mixture are evaluated by the `GlobalComponentEnumerator` to determine if they will participate in integrase reactions. Multiple different `GlobalComponentEnumerators` can be added to represent different integrases or other processes like splicing. If a new `DNA_construct` is generated by the `GlobalComponentEnumerator`, that information is saved in the `IntegraseSites` that participated in the reaction (in the case of integrase `GlobalComponentEnumerator`). Once the recursion depth is satisfied, remaining `DNA_constructs` undergo local component enumeration. As part of this process, species and reactions are generated, and `IntegraseSites` make use of these combinatorially bound species to create the proper reactions between species that integrases would allow.

favorable than intermolecular reactions. It may also be possible to harness this complexity, as we aim to do with the event recorder system presented in the previous chapter. In any case, the ability to easily generate these very complicated CRNs allows integrase-containing systems to be studied more carefully and "undesired" or "unexpected" integrase reactions may be more easily predicted and accounted for.

The following code snippet shows how to set up a integrase sites inside a `DNA_construct` and use an `Integrase_Enumerator`. The output CRN (upt to recursion depth 2) is shown in Figure 3D-E.

---

```
#Create two general DNA parts (these are illustrative and could be replaced by
     functional parts like promoters)
part1 = DNA_part("X")
part2 = DNA_part("Y")

#Create integrase sites
attP = IntegraseSite(name = "attP", site_type = "attP", integrase = "Bxb1")
attB = IntegraseSite(name = "attB", site_type = "attB", integrase = "Bxb1")

#Create the integrase rules attB + attP -> attL, attP + attB -> AttR
bxb1_rule = IntegraseRule("Bxb1",
     reactions={("attB","attP"):"attL",("attP","attB"):"attR"})

#Create a global component enumerator which uses the above rule to enumerate integrase
     reactions
bxb1 = Integrase_Enumerator("Bxb1", int_mechanisms={"Bxb1":bxb1_rule})

#Create a dictionary of parameters
parameters={"cooperativity":2,"kb":100, "ku":10, "ktx":.05, "ktl":.2,
     "kdeg":2,"kint":.05}

#Create two DNA_constructs (because there are no promoters, terminators, etc. they will
     not do local component enumeration)
p1 = DNA_construct([attP, part1, attB],circular=True)
p2 = DNA_construct([attP, part2, attB],circular=True)

#Create a Mixture with the global component enumerator and plasmid components
M = SimpleTxTlExtract(
    name = "txtl", components = [p1, p2],
    parameters = parameters, global_component_enumerators=[bxb1]
)

#Choose a recursion depth for compilation
recursion_depth = 2

#Compile the CRN using the specified recursion depth
#note: enumerated components can also be returned
CRN, enumerated_comps = M.compile_crn(return_enumerated_components = True,
     recursion_depth = recursion_depth)
```

#### 1.10 Flippable Promoter Global and Local Component Enumeration Example

Due to the fact that global component enumeration and local component enumeration occur sequentially, these two compilation steps can be used together. In the following example, we show the code behind the example in 3F-H where an `Integrase_Enumerator` is used to generate `DNA_constructs` due to integrase reactions which, in turn, use local component enumeration to work out the details of combinatorial binding and generate the appropriate `RNA_constructs`. All the `Components` enumerated this way use BioCRNpyler `Mechanisms` in order to compile the final CRN, resulting in the ability to rapidly generate and configure very complex models.

```
#Create DNA Parts
pconst = Promoter("Pr") #constitutive promoter
rbs = RBS("RBS") #regular RBS
bbx1_cds = CDS("Bxb1") #codes for the integrase Bxb1
rfp = CDS("RFP") #codes for RFP
T = Terminator("T")
gen_ori = Origin("genome") #put this Origin part in the genome so it is not considered
     for intermolecular reactions
```

```
gen_ori.attributes = ["no_inter", "genomic"] #this is what tells us that any dna that
     contains this part should not be considered for intermolecular reactions

#Create the integrase sites
attP = IntegraseSite(name = "attP", site_type = "attP",integrase="Bxb1")
attB = IntegraseSite(name = "attB", site_type = "attB",integrase="Bxb1")

#Create the integrase rule
bxb1_rule = IntegraseRule("Bxb1",
     reactions={("attB","attP"):"attL",("attP","attB"):"attR"})

#Create an integase enumerator
bxb1 = Integrase_Enumerator("Bxb1", int_mechanisms={"Bxb1":bxb1_mechanism})

#Create a DNA_construct. Here, parts are part of sublists which include their orientation
flip_construct = DNA_construct(
    [[T,"reverse"],[rfp,"reverse"],[rbs,"reverse"],[attP,"forward"],[pconst,"forward"],[attB,"reve
         [T], gen_ori]
)

#Integration rate
parameters={"kint":.01}

#Create a Mixture with the components and the global component enumerator
M = TxTlExtract(
    name = "e coli extract",
    components = [flip_construct], global_component_enumerators=[bxb1]
    parameter_file = "default_parameters.txt", parameters = parameters
)

#To set the initial condition
initial_concentration_dict = {
    flip_construct.get_species():0.0001
}

#Compile the CRN
#Note: recursion depth doesn't matter when only one integration event is possible
#Note: enumerated components can also be returned
CRN, enumerated_comps = M.compile_crn(return_enumerated_components = True,
     initial_concentration_dict = initial_concentration_dict)
```

### 2 Supplemental: Tables of Features

This section lists many of the different `Mixture`, `Component` and `Mechanism` classes available in BioCRNpyler. For more details about these classes and examples using many of them, check out the examples folder on GitHub.

### 3 Supplemental: Creating Custom BioCRNpyler Classes

BioCRNpyler is designed to be easily extendable so even non-computer scientists can add their own custom functionality. In this section we briefly show how to subclass core BioCRNpyler classes. For more details and examples, interested readers should look at the developer overview on our github.

**Table 1.** CRN Species Classes the BioCRNpyler Library

| Class | Data Structure | Description |
|---|---|---|
| `Species` | - | Species class with no structure. |
| `ComplexSpecies` | Set | Two or more species bound together. |
| `OrderedComplexSpecies` | Static List | Species bound together in a linear order. |
| `OrderedPolymerSpecies` | Editable List | Species in a non-branching polymer. |
| `PolymerConformation` | Hypergraph | Species and polymers bound together in a secondary structure. |

**Table 2.** Reaction Propensities in the BioCRNpyler Library

| Class | Function | Description |
|---|---|---|
| `Massaction` | deterministic: $k\prod_i S_i^{I_i}$ stochastic: $k\prod_i \frac{S_i!}{(S_i-I_i)!}$ | $I_i$ are the number of inputs of species $S_i$ |
| `HillPositive` | $k\frac{s^n}{K^n+s^n}$ | $s$ is a species $k$, $K$, and $n$ are parameters. |
| `HillNegative` | $k\frac{1}{1+(\frac{s}{K})^n}$ | $s$ is a species $k$, $K$, and $n$ are parameters. |
| `ProportionalHillPositive` | $kd\frac{s^n}{K^n+s^n}$ | $d$ and $s$ are species $k$, $K$, and $n$ are parameters. |
| `ProportionalHillNegative` | $kd\frac{1}{1+(\frac{s}{K})^n}$ | $d$ and $s$ are species $k$, $K$, and $n$ are parameters. |
| `GeneralPropensity` | $\rho(s)$ | $\rho$ can be written as a string. |

### 3.1 Mechanisms

Developing custom `Mechanisms` is also as easy as making a subclass of `Mechanism` and defining three functions to produce the desired CRN:

```
class CustomMechanism(Mechanism):
    def __init__(self, args, **kwargs ):
        Mechanism.__init__(self, name="name", mechanism_type="type", **kwargs)
        # python code to set internal variables

    def update_species(self, ... ):
        # python code to create Species objects
        return species_list

    def update_reactions(self, ... ):
        # python code to create Reaction objects
        return reaction_list
```

### 3.2 Components

It is also straightforward to make custom `Components`: simply subclass `Component` and define three functions:

```
class CustomComponent(Component):
    def __init__(self, args, **kwargs ):
        Component.__init__(self, ... , **kwargs)
        # python code to set internal variables
```

**Table 3.** Some `Mechanisms` in the BioCRNpyler Library

| Mechanisms Type | Mechanism Name | Description |
|---|---|---|
| binding | Reversible_Bimolecular_Binding | $S_1 + S_2 \leftrightharpoons (S_1 : S_2)$ |
| cooperative_binding | One_Step_Cooperative_Binding | $n\,S_1 + S_2 \leftrightharpoons (nS_1 : S_2)$ |
| cooperative_binding | Two_Step_Cooperative_Binding | $n\,S_1 \leftrightharpoons (nS_1)$, $(nS_1) + S_2 \leftrightharpoons (nS_1 : S_2)$ |
| cooperative_binding | Combinatorial_Cooperative_Binding | Allows a set of species $S_i$ and cooperativities $n_i$ to bind to a target $T$ in any order to form $(n_1 S_1 : ... : n_k S_k : T$ along with all combinatorial intermediaries. |
| binding | One_Step_Binding | $S1 + S2 ... SN \leftrightharpoons S1 : S2 : ... : SN$ |
| catalysis | BasicCatalysis | $S + C \rightarrow P + C$ |
| catalysis | BasicProduction | $C \rightarrow P + C$ |
| catalysis | MichaelisMenten | $Sub + Enz \leftrightharpoons Sub : Enz \rightarrow Enz + Prod$ |
| catalysis | MichaelisMentenReversible | $Sub + Enz \leftrightharpoons Sub : Enz \leftrightharpoons Enz : Prod \leftrightharpoons Enz + Prod$ |
| copy | MichaelisMentenCopy | $Sub + Enz \leftrightharpoons Sub : Enz \rightarrow Sub + Enz + Prod$ |
| transcription | OneStepGeneExpression | $G \rightarrow G + P$ |
| transcription | SimpleTranscription | $G \rightarrow G + T$ |
| translation | SimpleTranslation | $T \rightarrow T + P$ |
| transcription | PositiveHillTranscription | $G \rightarrow [r] G + P \;\; r = kG(R^n)/(K + R^n)$ |
| transcription | NegativeHillTranscription | $G \rightarrow [r] G + P \;\; r = kG/(K + R^n)$ |
| transcription | Transcription_MM | $G + RNAP \leftrightharpoons G : RNAP \rightarrow G + RNAP + mRNA$ |
| translation | Translation_MM | $mRNA + Rib \leftrightharpoons mRNA : Rib \rightarrow mRNA + Rib + Protein$ |
| transcription | multi_tx | $DNA : RNAp_n + RNAp \leftrightharpoons DNA : RNAp_n^{closed} \rightarrow DNA : RNAp_{n+1} DNA : RNAp_n \rightarrow DNA : RNAp_0 + nRNAp + nmRNA DNA : RNAp_n^{closed} \rightarrow DNA : RNAp_0^{closed} + nRNAp + nmRNA$ for $n = \{0, \max_{occ}\}$ |
| translation | multi_tl | $mRNA : RBZ_n + RBZ \leftrightharpoons mRNA : RBZ_n^{closed} \rightarrow mRNA : RBZ_{n+1} mRNA : RBZ_n \rightarrow mRNA : RBZ_0 + nRBZ + nProtein mRNA : RBZ_n^{closed} \rightarrow mRNA : RBZ_0^{closed} + nRBZ + nProtein$ for $n = \{0, \max_{occ}\}$ |
| dilution | Dilution | $s \rightarrow \emptyset$ |
| rna_degredation_mm | Degredation_mRNA_MM | $T + Nuclease \leftrightharpoons T : Nuclease \rightarrow Nuclease$. Global mechanism effects all RNA species $T$. ComplexSpecies containing RNA species are broken apart via the reaction $T : X + Nuclease \leftrightharpoons T : X : Nuclease \rightarrow X + Nuclease$. for any $X$. |
| degredation | Deg_Tagged_Degredation | $X + Protease \leftrightharpoons X : Protease \rightarrow Protease$. Here $X$ is any Species with the deg_tag attribute passed into the constructor of this GlobalMechanism. |

**Table 4.** Some `Components` in the BioCRNpyler Library

| Component Type | Component Name | Description |
|---|---|---|
| Chemical Complex | ChemicalComplex | A complex that represents the combination of several `Species`. Takes care of binding and unbinding reactions needed to form the complex |
| Chemical Complex | CombinatorialComplex | A complex that represents the combination of several `Species`. Binding occurs combinatorially in many possible orders. Allowed and disallowed intermediate complexes can be specified. |
| Enzyme | Enzyme | An enzyme that converts substrates to products |
| Protein | Protein | Basic component that represents a protein |
| DNA | DNA | Basic component that represents a DNA sequence |
| DNA | DNAassembly | A relatively simple DNA sequence containing one promoter, RBS, and a product |
| DNA | DNA_construct | A more complex DNA sequence that can have any number of `Components` in any order |
| Promoter | Promoter | Constitutive $\sigma 70$ promoter |
| Promoter | RegulatedPromoter | Repressible or activatable promoter such as $P_{tet}$ |
| Promoter | ActivatablePromoter | Activatable promoter using a positive Hill function |
| Promoter | RepressiblePromoter | Repressible promoter using a negative Hill function |
| Promoter | CombinatorialPromoter | Flexible promoter mechanism allowing various transcription factor binding configurations to allow or prevent transcription |
| Promoter | CombinatorialConformationPromoter | Enumerates the binding and unbinding events to represent a promoter with many regulators that can also form various secondary structures such as loops. |
| Ribosome Binding Site | RBS | Simple RBS using a translation mechanism |
| Coding Sequence | CDS | Protein coding part used for `DNA_construct`. Doesn't affect CRN |
| Terminator | Terminator | Transcriptional terminator used for `DNA_construct`. Doesn't affect CRN |
| RNA | RNA | Basic component that represents an RNA sequence |
| RNA | RNA_construct | A more complex RNA sequence that can have any number of `Components` in any order. Usually automatically generated by `DNA_construct` |
| Polymer Secondary Structure | CombinatorialConformation | Enumerates the binding and unbinding events to produce a `PolymerConformation` with different bound complexes and secondary structure. |

```python
def update_species(self):
    # python code calls mechanism.update_species( ... )
    return species_list
```

**Table 5.** Some Mixtures in the BioCRNpyler Library

| Mixture Name | Description |
|---|---|
| ExpressionExtract | A model for gene expression without machinery such as ribosomes, polymerases, etc. Here transcription and translation are lumped into one reaction: expression. |
| SimpleTxTlExtract | A model for transcription and translation in a cell-free extract without machinery such as ribosomes, polymerases, etc. RNA is degraded via a global mechanism. |
| TxTlExtract | A model for transcription and translation in a cell-free extract with machinery for ribosomes, polymerases, and endonucleases action. This model does not include any energy buffer. |
| EnergyTxTlExtract | Transcription and translation with ribosomes, polymerases, and endonucleases labelled as cellular machinery. Also includes a simple model of biochemical energy utilization involving NTPs, amino acids, and energy regeneration from a food source. Adapted from the model in [1]. |
| ExpressionDilutionMixture | A model for *in-vivo* gene expression without any machinery such as ribosomes, polymerases, etc. Transcription and translation are lumped into one reaction and a global mechanism is used to dilute all non-DNA species. |
| SimpleTxTlDilutionMixture | Mixture with continuous dilution for non-DNA species. Transcription (TX) and Translation (TL) are both modeled as catalytic with no cellular machinery. mRNA is also degraded via a separate reaction to represent endonucleases. |
| TxTlDilutionMixture | Transcription and translation with ribosomes, polymerases, and endonucleases labelled as cellular machinery. Also includes a background load which represents innate loading effects in the cell. Effects of loading on cell growth are not modeled. It has global dilution for non-DNA and non-machinery species. This model does not include any energy. |

```
    def update_reactions(self):
        # python code calls mechanism.update_reaction( ... )
        return reaction_list
```

### 3.3 Mixtures

Making custom Mixtures is also easy — it can be done via simple scripts by adding Components and Mechanisms to a Mixture object:

```
MyMixture=Mixture("customized mixture",
                  components=[List Components],
                  mechanisms=dict("mechanism_type":Mechanism))
```

The Mixture class can also be easily subclassed by rewriting the constructor:

```
class CustomMixture(Mixture):
    def __init__(self, args, **kwargs ):
        #python code to set up internal variables,
        # create Components, and default Mechanisms
        Mixture.__init__(self, mechanisms=dict("mechanism_type": Mechanism),
                         components=[List of Components], **kwargs)
```
